# A Lipidomics-Based Method to Eliminate Negative Urine Culture in General Population

**DOI:** 10.1101/2024.04.11.589091

**Authors:** Linda K Nartey, Abanoub Mikhael, Helena Pětrošová, Victor Yuen, Pamela Kibsey, Robert K Ernst, Michael X Chen, David R Goodlett

**Author notes:** Address correspondence to David R Goodlett, and Michael X Chen.

## Abstract

Urinary tract infections (UTIs) pose a significant challenge to human health. Accurate and timely detection remains pivotal for effective intervention. Current urine culture techniques, while essential, often encounter challenges where urinalysis yields positive results, but subsequent culture testing produces a negative result. This highlights potential discrepancies between the two methods and emphasizes the need for improved correlation in urinary tract infection (UTI) detection. Employing advanced lipidomics techniques, we deployed the fast lipid analysis technique (FLAT) on a clinical cohort suspected of having UTIs. Lipid fingerprinting by MALDI-TOF MS, directly from urine samples without *ex vivo* growth, correctly identified the common uropathogens within a 1-hour timeframe when compared to urine culture. FLAT analysis also identified urine samples without culturable pathogens (negative UTIs) with 99% agreement, whereas urinalysis showed 37% agreement with the gold standard urine culture. In 402 urine samples suspected for UTI from out-patients, FLAT assay rapidly ruled out negative urines without the need for culture in 77% of all cases. The potential impact of this innovative lipidomic-based approach extends beyond conventional diagnostic limitations, offering new avenues for early detection and targeted management of urinary tract infections. This research marks a paradigm shift in urine culture methodology, paving the way for improved clinical outcomes and public health interventions.

## INTRODUCTION

Urinary tract infections (UTIs) are one of the most common diseases worldwide, affecting approximately 150 million people every year, resulting in approximately 3.5 billion US dollars in healthcare costs (1). As a result, suspected UTIs are one of the most common presentations in a primary care setting. Although several urine biomarkers have been investigated for the diagnosis of UTIs, urine culture remains the gold standard (2). Approximately 80% of requested urine cultures come from outpatient settings, with as many as 70–80% of these samples showing no significant uropathogens (3). These negative urine cultures generate a considerable workload, which occupies microbiology laboratory staff leading to a waste of laboratory resources. Additionally, empiric antibiotic treatments may potentially have no clinical benefit and may increase the risk of adverse events, antimicrobial resistance and healthcare costs.

Many patients with uncomplicated UTIs present clinically as straightforward cases that may not require additional testing beyond urinalysis. Because of the time-consuming nature of urine culture, many clinical laboratories have instituted reflexive workflows, such as urine culture, only after dipstick analysis has demonstrated positive findings. However, urine dipstick analysis for blood, nitrites, and leukocyte esterase can be prone to positive interferences leading to unnecessary urine culture investigations (4). Hence, it is clinically and operationally beneficial to develop a rapid alternative urine screening test in a culture-free manner with minimal human intervention.

Currently, matrix-assisted laser desorption/ionization time-of-flight (MALDI-TOF) mass spectrometry (MS) protein fingerprinting tests, such as the Bruker Biotyper, are standard microbial identification (ID) methods (5). These methods involve the use of MALDI-TOF MS to analyze the protein profiles of bacterial samples selected after culture. By comparing detected protein profiles to previously recorded protein profiles for various bacterial species in a library, bacteria can be identified. However, the protein-based Biotyper method requires microbial growth to produce a pure colony that often requires at least 18-48 hours to produce a pure colony. Here, we use a similar approach to identify bacteria by MALDI-TOF MS analysis, but instead of protein profiles, we use lipids (6). Complex and diverse lipids are a major component of bacterial membranes. In Gram-positive bacteria, the cell envelope consists of complex multi-layers of peptidoglycan enclosing a single membrane, a bilayer of lipids including cardiolipin (CL) and lipoteichoic acid (LTA) (7). In contrast, Gram-negative bacteria have a two-bilayer membrane with the inner membrane consisting of a single layer of peptidoglycan. Lipid A, the endotoxic portion of lipopolysaccharide (LPS), is embedded in the outer membrane of Gram-negative bacteria and exhibits species-specific structural diversity (7). These bacterial cell wall lipid components have been shown to provide a unique lipid barcode for the identification of individual microbial species in a manner similar to protein fingerprinting (6). These lipid barcodes for each bacteria represent signature ions allowing individual bacteria to be uniquely identified. The microbial lipid biomarkers used in this study consist mainly of cardiolipin and Lipid A.

A recent study by Sorenson et al. (8) presented a method for direct lipid extraction on a MALDI plate, referred to as Fast Lipid Analysis Technique (FLAT). Similar to the chemistry used in traditional lipid microextraction, FLAT differs mainly in extraction of the aforementioned microbial lipids directly onto a MALDI plate. Thus, the FLAT assay reduces the effort needed to observe the lipids that serve as barcodes to identify bacteria and fungi, as well as detect membrane-based antimicrobial resistance (6,9–11). Given that FLAT circumvents the need for culture, results can be ready within an hour of the lab receiving a sample. Additionally, FLAT uses clinic-friendly reagents and has low hands-on time. Previously, Yang et al. have shown that microbial lipid profiles can accurately identify microbial colonies directly from known Gram-negative urine samples via FLAT with MALDI-TOF MS (6,9). However, their study only focused on analysis of known Gram-negative UTI urine samples.

Following the work of Yang et al., we used lipid biomarkers analyzed by MALDI-TOF MS to determine FLAT’s ability to discern positive from negative urines in a large clinical cohort. Over 400 suspected UTI samples from outpatient settings, excluding pregnant women and sexually transmitted disease (STD) patients, were used for this study. The overall aim of this study was to evaluate the use of FLAT to eliminate the need for urine culture in the general population.

## MATERIALS AND METHODS

### Ethics statement

This study was approved by the Vancouver Island Health Authority (VIHA) REB # H21-03785. All research took place at the core laboratory of Victoria General Hospital in Victoria, British Columbia.

### Sample collection

In this study, 435 urine specimens were collected from outpatients suspected to have UTIs from Victoria General and Royal Jubilee Hospital, British Columbia. Maternity care and sexually transmitted infection (STI) clinics were excluded. Urine samples were requested by clinicians as part of urinary tract infection investigations, either by 1) urinalysis (dipstick - Roche diagnostics, 11063 Stn CV, Montreal, QC) and urine culture or 2) urinalysis with reflex to urine culture. The criteria for reflex to urine culture were established based on positive urinalysis results incorporating any single or combined indications, including positive hemoglobin (Hb), nitrites, and leukocyte esterase. Urine cultures were identified using the protein Biotyper (MBT Sirius, Bruker, Bremen, Germany). Based on the inclusion criteria, all urine samples were collected from morning midstream clean-catch urine, without considering age and gender of the patients. Urine culture was not performed on 33 of the 435 (7.5%) samples that did not pass inclusion criteria.

The remaining 402 urine samples were tested by urinalysis and urine culture as part of the VIHA standard of care. The urine samples were obtained in sterile BD Vacutainer^®^ urine C&S Preservative Tubes (BD, 1 Becton Drive, Franklin Lakes. NJ). Specifically, a 1 mL aliquot was allocated from each sample for FLAT analysis within 3 days of receipt of the UTI sample. FLAT results were compared with corresponding urinalysis and urine culture reports. The VIHA microbiology laboratory does not perform further UTI investigations in urine cultures reported as “light growth”, “mixed growth”, “no growth” and “no uropathogens isolated”. Hence, all such reported outcomes were classified as negative urine cultures in this study.

### FLAT Analysis

Lipid A mass spectral analyses were performed using the FLAT assay described by Sorensen et al (8). A 1 mL aliquot of each urine was transferred into an Eppendorf tube and analyzed using FLAT.

For this study, stainless-steel, 96-well, disposable MALDI target plates (MFX μFocus plate 12×8 c 2,600 μm 0.7 T; Hudson Surface Technology, Inc., South Korea) were used. Urine samples were processed two ways. First, 1 µL of urine was spotted directly onto the MALDI target plate, and second, 1 mL of urine was centrifuged at 8,000g for 5 min to generate a urine pellet, after which 1 µL bacterial pellet was spotted onto a MALDI target plate. Next, 1 µL of a citric acid buffer containing 0.2 M anhydrous citric acid and 0.1 M trisodium citrate dihydrate (Fisher Chemical) was placed on top of each dried sample. The MALDI target plate was then incubated at 110°C for 30 min in a humidified chamber and gently rinsed with endotoxin free water. Each sample was spotted on the MALDI plate in triplicate. Subsequently, 1µL norharmane (10mg/mL) matrix (Sigma Aldrich) dissolved in 2:1 v/v MS-grade chloroform and methanol (both from Fisher Chemical) was spotted onto the extracted lipid sample on a MALDI target plate, allowed to dry and then analyzed by MALDI-TOF MS.

### Modifications to FLAT

Cloudy samples were diluted in 10-fold prior to FLAT analysis. Briefly, 100 µL of cloudy urine was diluted in 900 µL sterile deionized water. Next, 1 µL of diluted urine was spotted directly onto the MALDI target and analyzed by FLAT assay described above.

A second cohort of 70 known Gram-positive urine samples were analyzed by adding a 10 minute sonication before FLAT analysis. Briefly, 1 mL of urine was sonicated for 10 mins, 1 µL of urine was spotted directly onto the MALDI target plate, and then centrifuged at 8,000g for 5 min to generate a urine pellet, after which 1 µL bacterial pellet was spotted onto a MALDI target plate. Acidified and incubated in a humidified chamber at 110°C for 30 min. The plate was rinsed and 1 µL of Norharmane matrix was added to each spot. Samples were then analyzed by MALDI-TOF MS

### MALDI-TOF MS analysis and pathogen identification

MALDI-TOF MS analysis was conducted using a Bruker Microflex (Bruker, Bremen, Germany) in negative ion and linear mode. Analyses were conducted at 65% global intensity with 500 laser shots for each acquired mass spectrum. Mass spectra were collected between 1000 *m/z* to 2000 *m/z*. Acquired mass spectral data were processed using flexAnalysis (v3.4) software with smoothed and baseline corrections using monophosphoryl Lipid A (MPLA) as a post-acquisition mass calibrant. After MALDI-TOF MS analysis, pathogens were identified by comparing sample mass spectra to a developed microbial library (SI 1) and previously acquired reference spectra (6,8,9,12). In routine standard of care, cultures were identified using the protein Biotyper (MBT Sirius, Bruker, Bremen, Germany) and the 3.4.207.48. /BDAL 11 organism library.

### Identification of non-pathogenic markers by FLAT

Analyses of molecules of unknown structure at specific *m/z* values, some of which interfered with known microbial lipid signature ions, were analyzed using FLAT*^n^* (11), the tandem-MS version of FLAT on a Bruker MALDI Trapped Ion Mobility Spectrometry (TIMS) TOF MS (Bruker, Bremen, Germany). The instrument was calibrated before every experiment by direct infusion of electrospray tuning mix (Agilent, Santa Clara, CA, USA). To acquire tandem mass spectra, the precursor ion *m/z* value of interest was entered to the third decimal point with isolation width and collision energy set to 4 *m/z* and 100 – 110 eV, respectively. Mass spectra were acquired in negative ion mode with a mass resolution of 60,000 at *m/z* 400. After manual analysis of the tandem mass spectra of unknown ions, structures were hypothesized using mMass (v5.5.0) software and subsequently confirmed by manual analysis in ChemDraw v18.0 (PerkinElmer informatics, Waltham, MA, USA).

### Analysis of known Gram-positive cohort

The FLAT assay was performed on a second cohort of 70 known Gram-positive urine samples with slight modification. Briefly, 1 mL urine was sonicated for 10 minutes before analyzing via FLAT on both direct urine and pellets from this modification.

### Data analysis

Specificity and sensitivity of the results were calculated using the following formulas: SE=TP/(TP+FN) ×100% and SP=TN/(TN+FP) ×100% where the positive predictive value (PPV) and negative predictive value (NPV) were calculated as follows: PPV=TP/P; NPV=TN/N where FN = false negative; FP = false positive; N = negative; P = positive; SE = sensitivity; SP = specificity; TN = true negative; and TP = true positive). Mass spectral data were analyzed using flexAnalysis (v3.4) software processed with smoothed and baseline corrections.

## RESULTS

A total of 435 samples were collected for this study (Fig 1, 2). Of those, 33 coming from maternity care or STI clinics were excluded leaving 402 urine samples for analysis. The age of the study population ranged from 15 months to 98 years. Out of the 402 samples, 95% were aged >18 years old (42% were >65), 56% were female, and one patient was identified as gender X (Table 1). Overall, urinalysis reported 71% and 29% of the samples as positive and negative, respectively; whereas urine culture reported 22% and 78% of the samples as positive and negative. (Table 2 and Fig 3). The majority of positive urine cultures showed Gram-negative uropathogens (78%), with *Escherichia coli* being the most common at 78% (Table 2).

**Fig 1:**
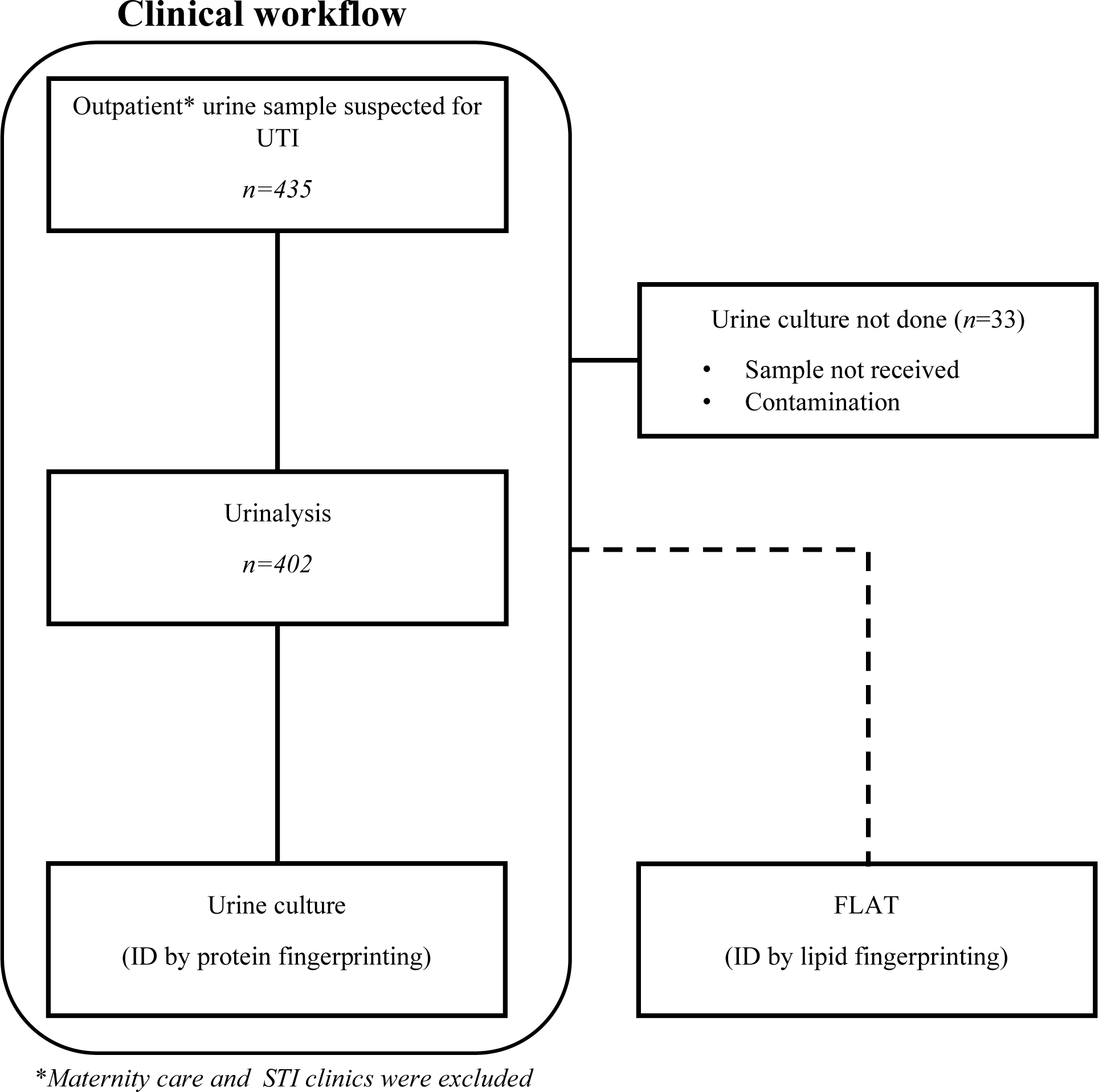
Study cohort inclusion/exclusion criteria.

**Fig 2:**
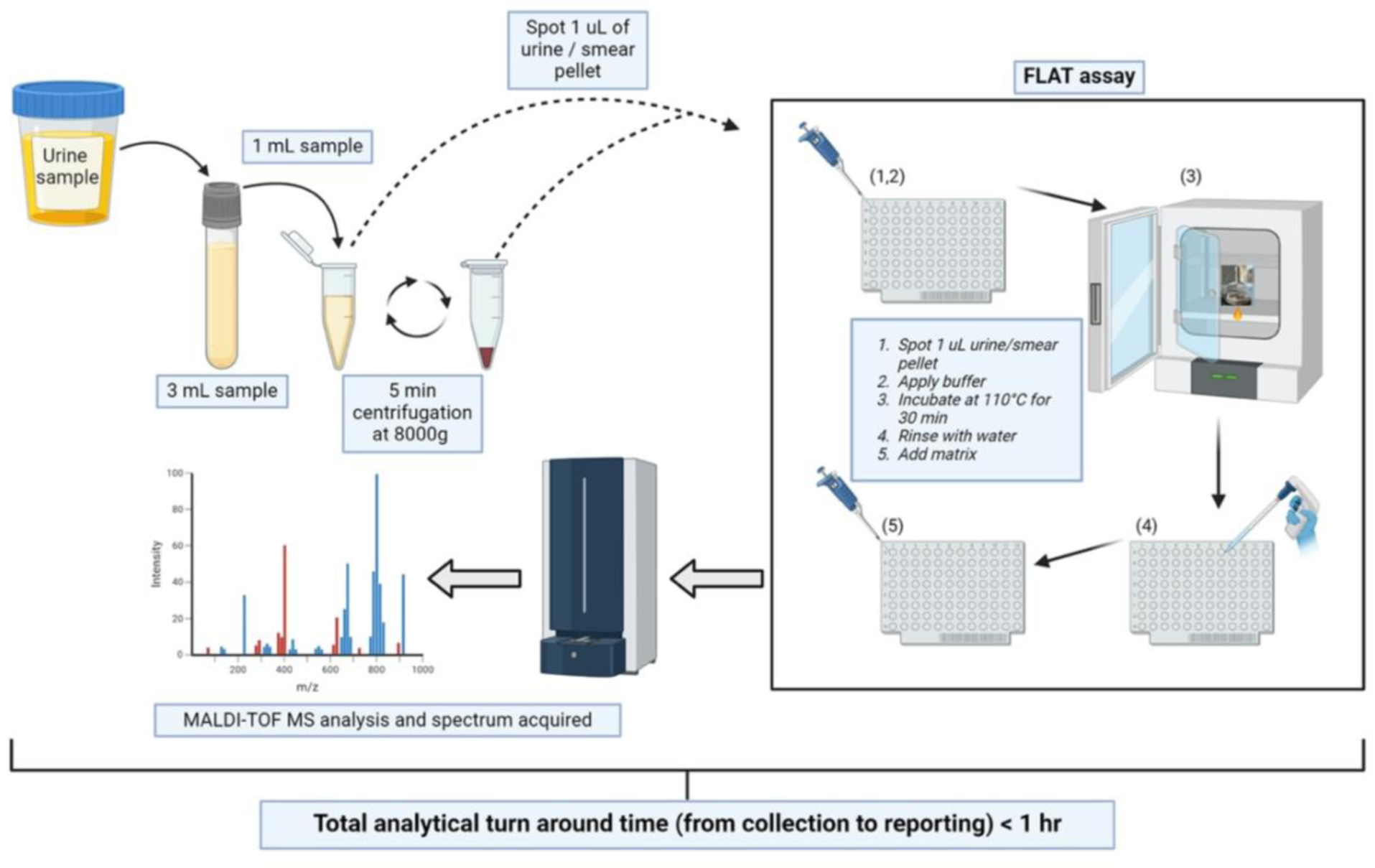
Experimental workflow for direct-from-urine analysis. Table 1: Patients demographics (n=402)

**Table 1:** Patients demographics (n=402)

|  | FEMALE | MALE | GENDER X |
| --- | --- | --- | --- |
| Children (1-12) | 10 | 4 |  |
| Adolescents (13-17) | 2 | 4 |  |
| Adults (18 or older) | 110 | 59 | 1 |
| Older adults (>65) | 103 | 109 |  |

**Fig 3:**
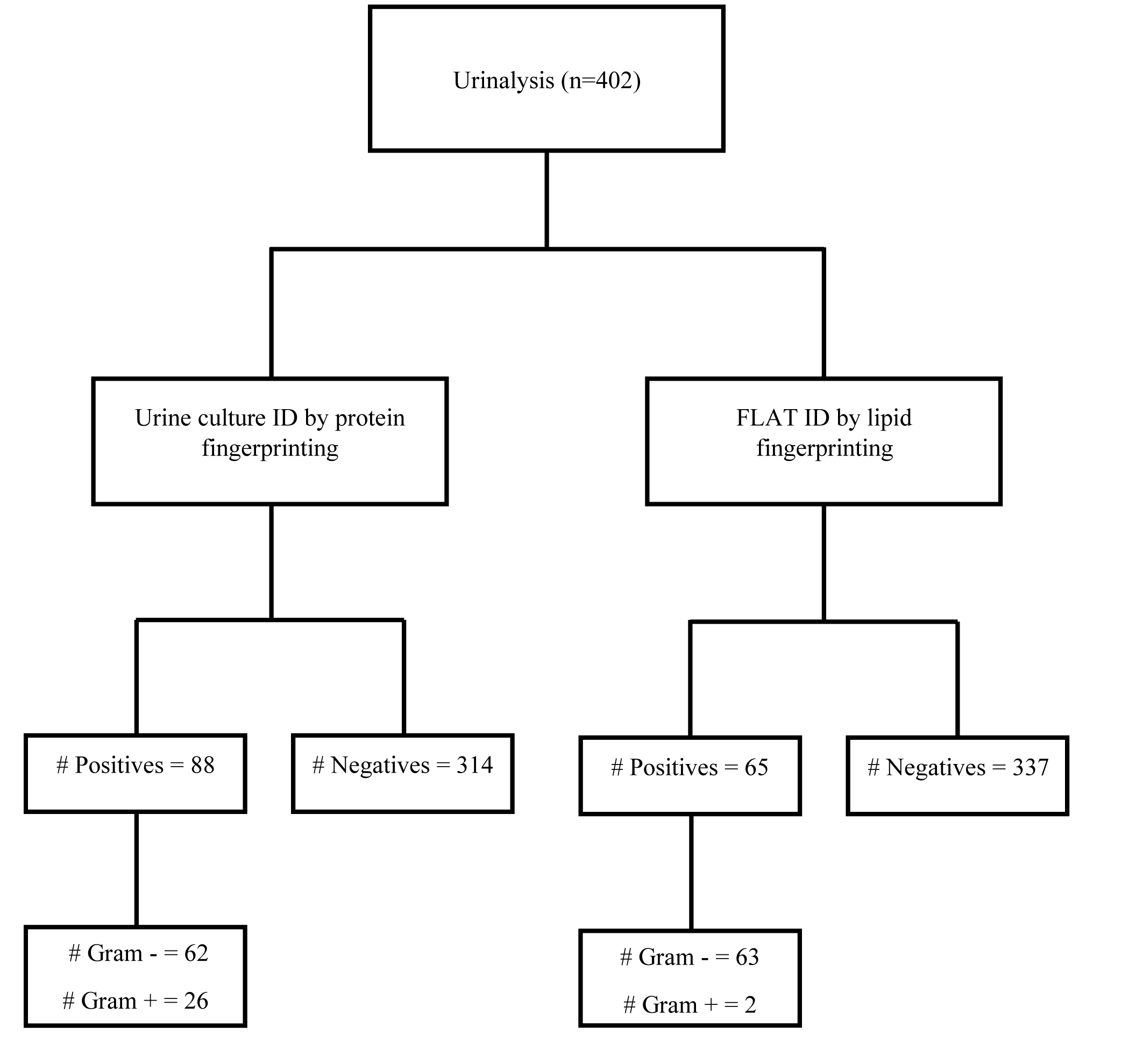
Comparison of pathogen identification by proteins and lipids.

**Table 2:** Positivity of urine culture and urinalysis in the study cohort.

| *n* = 402 | POSITIVE | NEGATIVE |
| --- | --- | --- |
| **CULTURE** | 88(22%) | 314 (78%) |
| **URINALYSIS** | 285(71%) | 117 (29%) |

### Modifications to FLAT

Approximately ∼3% of the 402 samples were observed to be cloudy containing sediment, resulting in noisy MALDI-MS profiles. To overcome this problem, these cloudy samples were diluted 10 fold, after which negative samples produced less noisy MALDI-MS spectra whereas positive samples (0.7%) produced mass spectra with high signal to noise (S/N) ratio bacterial lipid ions. Furthermore, only 8% of the 402 samples known to be Gram positive were detected by FLAT. To improve detection of Gram positives, sonication prior to FLAT was incorporated. In an analysis of a second cohort of 70 known Gram positive samples, this improved detection of Gram positives to 51%.

### Identification of uropathogens by FLAT

Directly from 1 µL of urine, we identified common uropathogens including fifty-one *Escherichia coli* (78%), six *Klebsiella* spp (9%), one *Pseudomonas aeruginosa* (2%), three *Enterobacter cloacae* (5%), three *Proteus mirabilis* (5%), and one sample that was polymicrobial containing *P. aeruginosa* and *E. coli* (Fig 4). Two Gram-positive bacteria, *S. epidermidis* and *A. urinae* were also detected (Fig 5).

**Fig 4:**
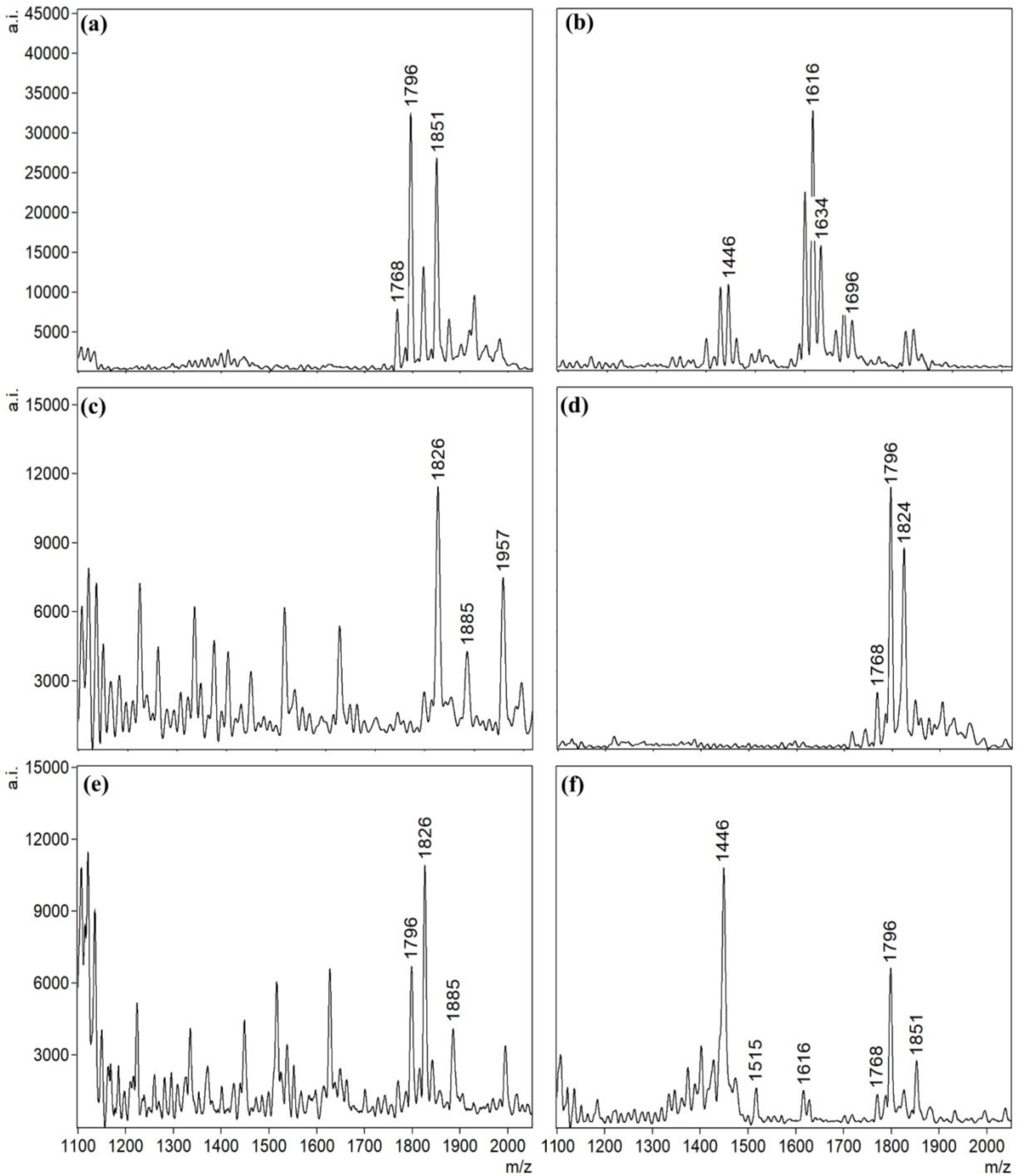
Representative FLAT mass spectra of detected Gram-negative uropathogens. (a) *E. coli,* (b) *P. aeruginosa,* (c) *Proteus* sp, (d) *Enterobactor sp,* (e) *Klebsiella* sp, and (f) Poly-microbial (*P. aeruginosa* and *E. coli*)

**Fig 5:**
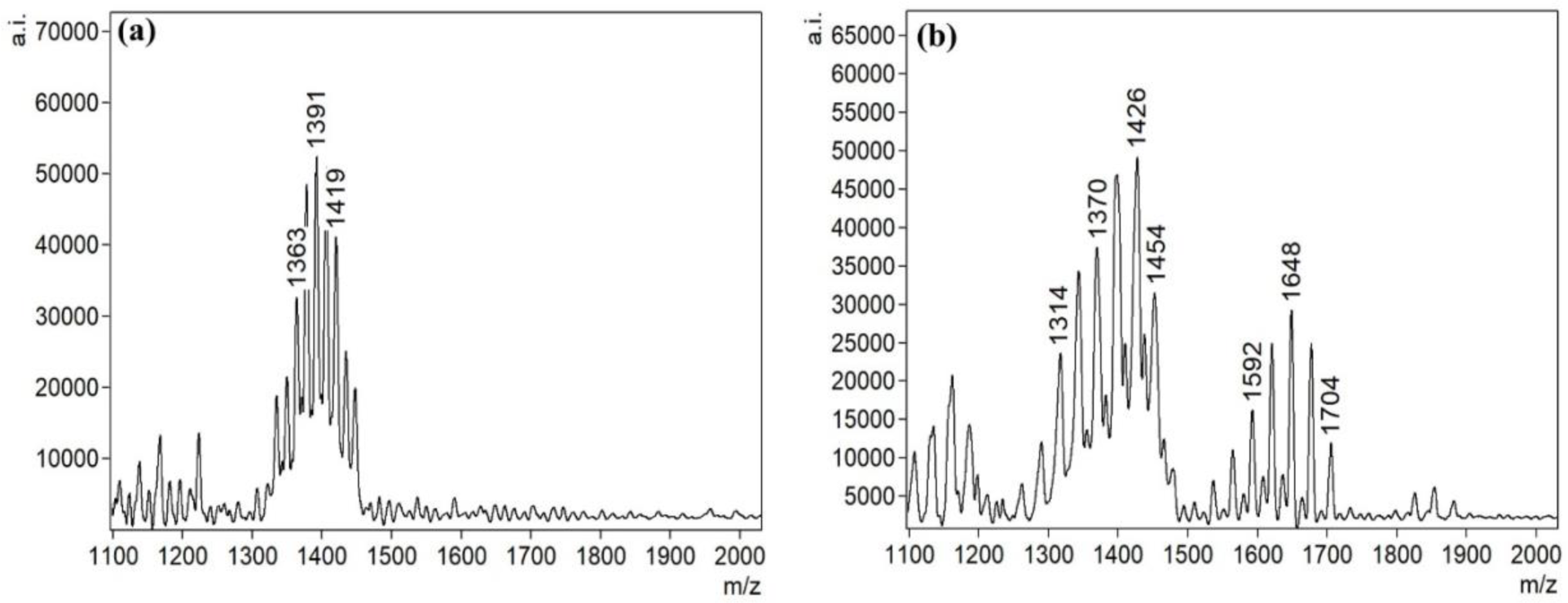
Representative FLAT mass spectra of detected Gram-positive uropathogens. (a) *S. epidermidis* and (b) *A. urinae*.

### Method comparison

The FLAT and culture results showed 93% concordance (Table 3). Overall, the FLAT assay had a sensitivity of 70% and specificity of 99% with positive and negative predictive values of 93 and 99%, respectively (Table 4). For negative urine cultures the FLAT assay was 99% in concordance whereas urinalysis was 37%. (Table 4a). Additionally, the FLAT assay was 97% (n=88) and 8% (n=26) in agreement with Gram-negative and Gram-positive cultures, respectively (Table 4b). An additional 70 known Gram-positive urines were obtained to increase the sample size. After adding a 10 min sonication step prior to FLAT, Gram-positive bacterial detection increased from 8% of the initial cohort of 402 to 51% of the new cohort of 70 samples.(Table 5).

**Table 3:** A 2 x 2 diagnostic table of FLAT assay.

|  |  | UTI Results |  |  |
| --- | --- | --- | --- | --- |
|  |  | Culture + | Culture - |  |
| FLAT Results | Positive | True Positive<br>(62) | False Positive<br>(3) | Positive predictive<br>value = 95% |
|  | Negative | False Negative<br>(26) | True Negative<br>(311) | Negative predictive<br>value = 92% |
|  |  | Sensitivity = 70% | Specificity = 99% |  |

**Table 4:** Method comparison of FLAT, urinalysis and culture: (a) Result comparison of negative urines (b) Result comparison of positive urines.

| (a) CULTURE | URINALYSIS (dipstick) | FLAT | COMMENTS |
| --- | --- | --- | --- |
| NEG | NEG | NEG |  |
| N= 314 | 117 (37%) | 311 (99%) | FLAT Oversensitive (3 false positives by FLAT analysis) |

| (b) CULTURE | URINALYSIS | FLAT POS | COMMENTS |
| --- | --- | --- | --- |
| POS | (DIPSTICK) POS |  |  |
| **Gram –**<br>**(n=62)** | 62 (100%) | 60 (97%) | 2 undetected bacteria by FLAT<br>(Acinetobacter and Citrobacter<br>spp) |
| **Gram +**<br>**(n=26)** | 24 (92%) | 2 (8%) |  |

**Table 5:**
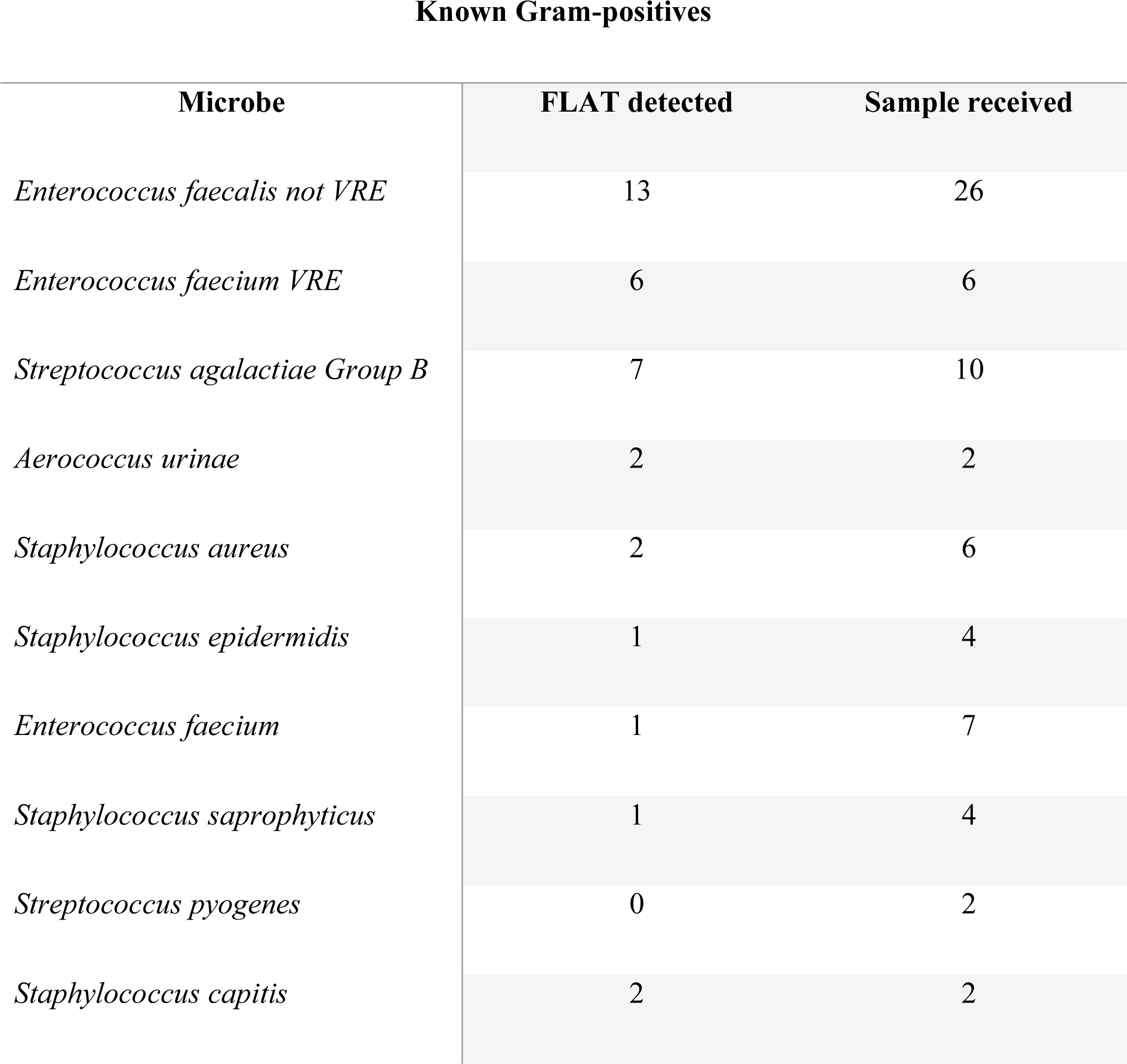
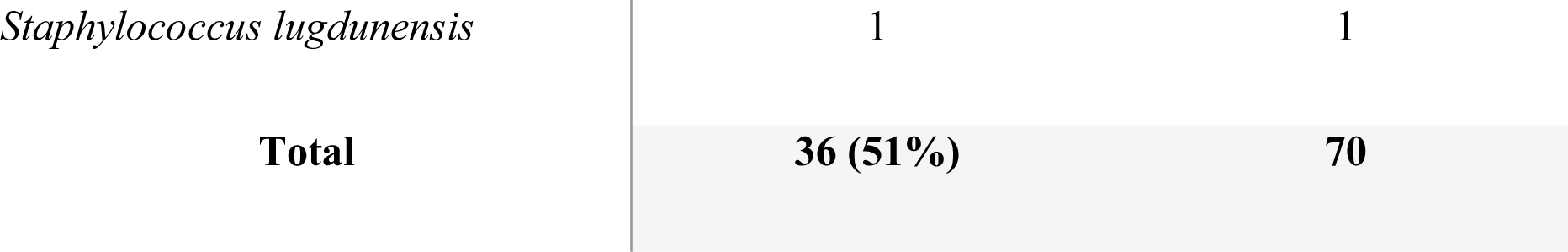
Detection of Gram-positive bacteria by FLAT.

### Identification of non-pathogenic urine components by FLAT

In addition to microbial lipids, FLAT detected host cardiolipins and heme degradation products. Approximately 40% of urine samples showed a cluster of ions at *m/z* 1447-1449, which is similar to one of the signature ion of *P. aeruginosa* Lipid A (Fig 4b). However, samples containing these ions of *m/z* ∼1447-1449, by FLAT failed to produce viable colonies when isolated on a culture media (Fig 6b). Tandem MS results of the *m/z* 1448, ion cluster showed similar fragmentation to a mammalian cardiolipin reported by Kim et al. (13) (SI. Fig 1).

**Fig 6:**
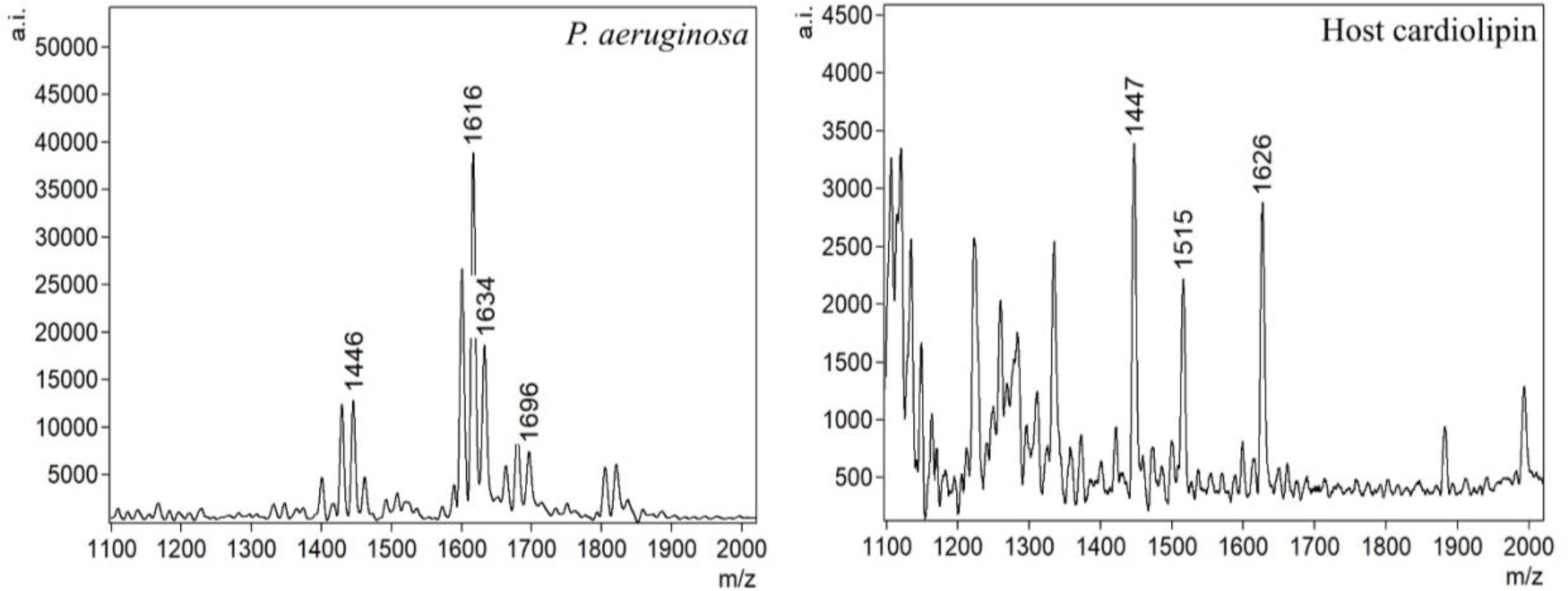
FLAT mass spectra showing *m/z* 1447-1449 interference (a) Lipid A mass spectra of *P. aeruginosa* and (b) Host cardiolipin.

Additionally, an ion at *m/z* 1230, which was present in all urine samples with blood (6.5% of initial cohort of 402) (Fig 7a), was detected by FLAT. After FLAT analysis on a plasma Hb standard obtained from the clinical laboratory, we observed the presence of a similar ion at *m/z* 1230 (Fig 7b). Manual analysis of the tandem MS spectrum of this ion compared against the same *m/z* value from the plasma standard predicted this ion to be a heme dimer (SI Fig 2).

**Fig 7:**
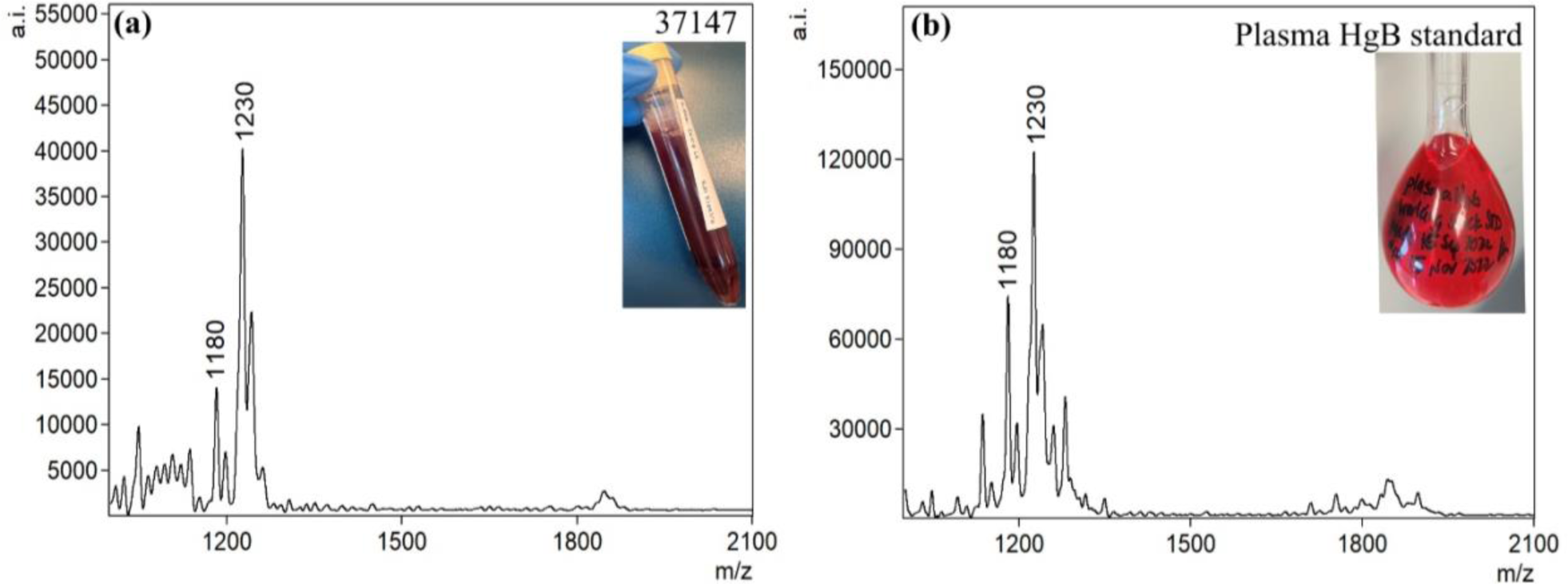
FLAT mass spectra of heme: (a) Patient urine with blood (b) Plasma HgB standard.

## DISCUSSION

The culture of urine samples remains the gold standard for diagnosing UTIs, but this process takes 24–72 hours to complete, and inappropriate sampling or storage conditions can lead to false positive results (14). A rapid and inexpensive test with high negative predictive value would reduce unnecessary urine culture and inappropriate empiric antimicrobial treatment. In this study, we present FLAT, a lipid-based method that allows direct lipid extraction on a MALDI plate for pathogen detection in under an hour from receipt of a sample. We investigated the diagnostic performance of FLAT in a clinical cohort to highlight the potential benefits of this assay in comparison with conventional urine culture.

FLAT analysis identified urine samples without culturable pathogens (negative UTIs) with 99% agreement, whereas urinalysis showed 37% agreement with the gold standard urine culture. This technique shows that microbial membrane lipids analyzed by MS represent biomarkers that can be used to identify mono- or poly-microbial specimens directly from urine samples without the need for culture. The FLAT assay identified common uropathogens directly from urine samples within 1-hour of collection. In 402 urine samples suspected for UTI from out-patients, FLAT assay rapidly ruled out negative urines without the need for culture in 77% of all cases.

It is estimated that over 80% of UTIs are caused by Gram-negative bacteria and predominantly by *E. coli* (15). From our study, the FLAT data showed an overall sensitivity of 70% and specificity of 99% with positive and negative predictive values of 95 and 92%, respectively for both Gram-positive and -negative bacteria. The low sensitivity is the result of inconsistent detection of Gram-positive bacteria, which might be due to relatively lower amounts of cardiolipin signature ions being released from Gram positives than lipid A from Gram negatives in these urine samples (16,17). Additionally, the FLAT extraction process may be less effective in disrupting Gram-positive cell walls. Given that FLAT was developed to isolate lipid A from Gram-negative bacteria, it is likely that the rigidity of the Gram-positive cell wall prevents the efficient release of cardiolipin from Gram-positive bacteria. This seems likely given that lipids in general have good ionization efficiency (18) and is bolstered by the report of Angelini et al who showed the use of cardiolipin fingerprinting by MALDI-TOF as a screening tool with no difficulties in ionization (19).

To improve the sensitivity of FLAT, centrifugation of urine samples prior to analysis was carried out in an attempt to enrich pathogens. Gram-negatives showed an improved sensitivity of 94% and specificity of 99% with positive and negative predictive values of 95 and 99%, respectively. In contrast, centrifugation did not improve detection of Gram-positive bacteria. (15) To overcome this diagnostic limitation, a 10 minutes sonication step was added to sample preparation. It was also noted less than 7% of samples tested positive for Gram-positive bacteria in our cohort of 402 samples. Subsequently, we obtained another 70 urine samples from the microbiology lab with confirmed growth of Gram-positive bacteria. Improved cardiolipin detection was observed after sonication with an increased sensitivity for Gram-positive bacteria to 51%. We believe this increased cell wall disruption allowed for a more comprehensive release of the cardiolipins leading to a higher detection rate. This implies that, had sonication been used in the original cohort, sensitivity of FLAT would have been about 85% due to increased detection of Gram-positive containing samples. Encouraged by this outcome, we are continuing method development to further improve Gram positive detection.

The mass range of bacterial cardiolipin and lipid A is between *m/z* 1000-2000. In some cases, there were interfering ions present that initially led to false positives. To prevent such false positive identifications, it was essential to define non-microbial ions present in this mass range of interest that were present at similar *m/z* values to the signature ions (see supplemental Table 1). Urine, being a by-product of metabolism, can contain non-microbial chemical components that might interfere with FLAT identification of pathogens via their signature ions. For example, in this study, approximately 40% of the samples diagnosed as negative for bacteria showed an ion at *m/z* 1447-1449, which is similar to the reported as one of the signature ion for *P. aeruginosa* lipid A (6). However, tandem MS results of *m/z* 1448 showed similar fragments to what Kim et al. (13) reported as a mammalian cardiolipin. We note that this putative cardiolipin was found in suspected UTI samples that were both positive and negative for bacteria and were not gender or age specific.

In all urine samples positive for Hb, FLAT detected an ion at *m/z* 1230. With hematuria being one of the symptoms of UTIs, it was paramount to identify and characterize this ion. Tandem MS results of this ion compared against a plasma standard analyzed by FLAT confirmed it was, in fact, a heme dimer. In 1995, Benesch and Kwong reported that hemoglobin can easily be transferred between body fluids in the form of dimers (21). This phenomenon explains why MS did not detect heme *m/z* 614 but rather showed a heme dimer at *m/z* 1230.

There was one case of polymicrobial infection which was accurately identified by FLAT as *E. coli* and *P. aeruginosa* with distinct ions and their respective m/z were detected without interference. The ability to detect polymicrobial infections directly from urine within 1 hour will guide appropriate antibiotic therapy and improve clinical outcome (22).

## CONCLUSION

The ability to identify pathogens without need for culture can allow for faster pathogen identification, reduced time to appropriate antimicrobial therapy, reduced use of disposable plasticware and improved patient outcomes. Rapid exclusion of urinary tract infections and the elimination of culture-negative urines in low-risk population avoid unnecessary empiric antibiotic treatment and save healthcare resources. When compared to the current standard of care, FLAT demonstrated accurate microbial identifications of Gram-negative bacteria with an analytical turn-around-time of < 1 hour. The FLAT assay provides a rapid alternative urine screening test that out-performed urinalysis (dipstick) with lower false positive rate. Notably, FLAT was able to rule out UTI with 99% accuracy. Finally, we suggest that use of FLAT as a screening test for suspected UTIs could eliminate ∼77% of negative urine cultures in outpatient population within 1 hour, greatly reducing microbiology laboratory workloads by eliminating unnecessary culturing.

## Acknowledgments

Profs Goodlett and Ernst thank the National Institutes of Health for funding from R01AI147314. Dr Chen thanks the Vancouver Island Health Authority and Victoria hospitals foundation for funding from the Catalyst grant.

## SUPPLEMENTAL INFORMATION (SI)

**SI. Table 1:**
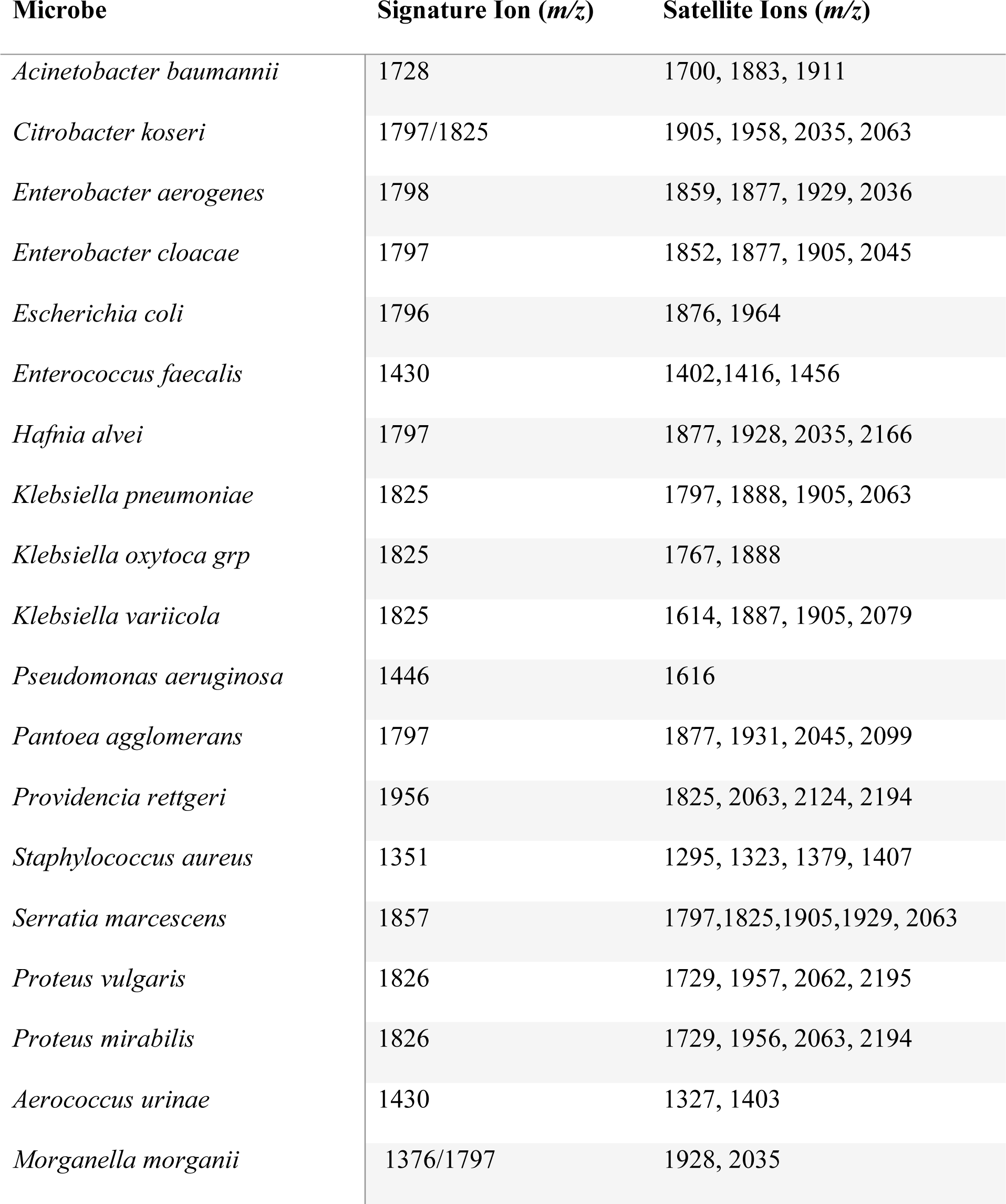
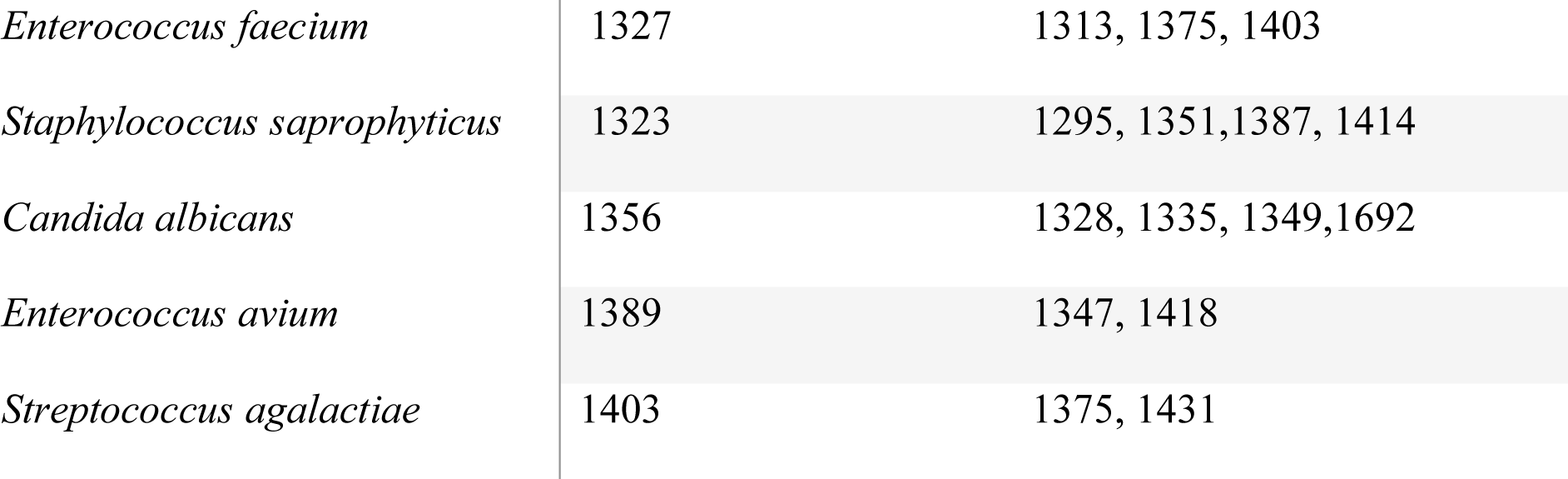
Microbial library of common uropathogens from study cohort.

| Microbe | Signature Ion (*m/z*) | Satellite Ions (*m/z*) |
| --- | --- | --- |
| *Acinetobacter baumannii* | 1728 | 1700, 1883, 1911 |
| *Citrobacter koseri* | 1797/1825 | 1905, 1958, 2035, 2063 |
| *Enterobacter aerogenes* | 1798 | 1859, 1877, 1929, 2036 |
| *Enterobacter cloacae* | 1797 | 1852, 1877, 1905, 2045 |
| *Escherichia coli* | 1796 | 1876, 1964 |
| *Enterococcus faecalis* | 1430 | 1402, 1416, 1456 |
| *Hafnia alvei* | 1797 | 1877, 1928, 2035, 2166 |
| *Klebsiella pneumoniae* | 1825 | 1797, 1888, 1905, 2063 |
| *Klebsiella oxytoca* grp | 1825 | 1767, 1888 |
| *Klebsiella variicola* | 1825 | 1614, 1887, 1905, 2079 |
| *Pseudomonas aeruginosa* | 1446 | 1616 |
| *Pantoea agglomerans* | 1797 | 1877, 1931, 2045, 2099 |
| *Providencia rettgeri* | 1956 | 1825, 2063, 2124, 2194 |
| *Staphylococcus aureus* | 1351 | 1295, 1323, 1379, 1407 |
| *Serratia marcescens* | 1857 | 1797, 1825, 1905, 1929, 2063 |
| *Proteus vulgaris* | 1826 | 1729, 1957, 2062, 2195 |
| *Proteus mirabilis* | 1826 | 1729, 1956, 2063, 2194 |
| *Aerococcus urinae* | 1430 | 1327, 1403 |
| *Morganella morganii* | 1376/1797 | 1928, 2035 |
| *Enterococcus faecium* | 1327 | 1313, 1375, 1403 |
| *Staphylococcus saprophyticus* | 1323 | 1295, 1351, 1387, 1414 |
| *Candida albicans* | 1356 | 1328, 1335, 1349, 1692 |
| *Enterococcus avium* | 1389 | 1347, 1418 |
| *Streptococcus agalactiae* | 1403 | 1375, 1431 |

**SI. Fig 1:**
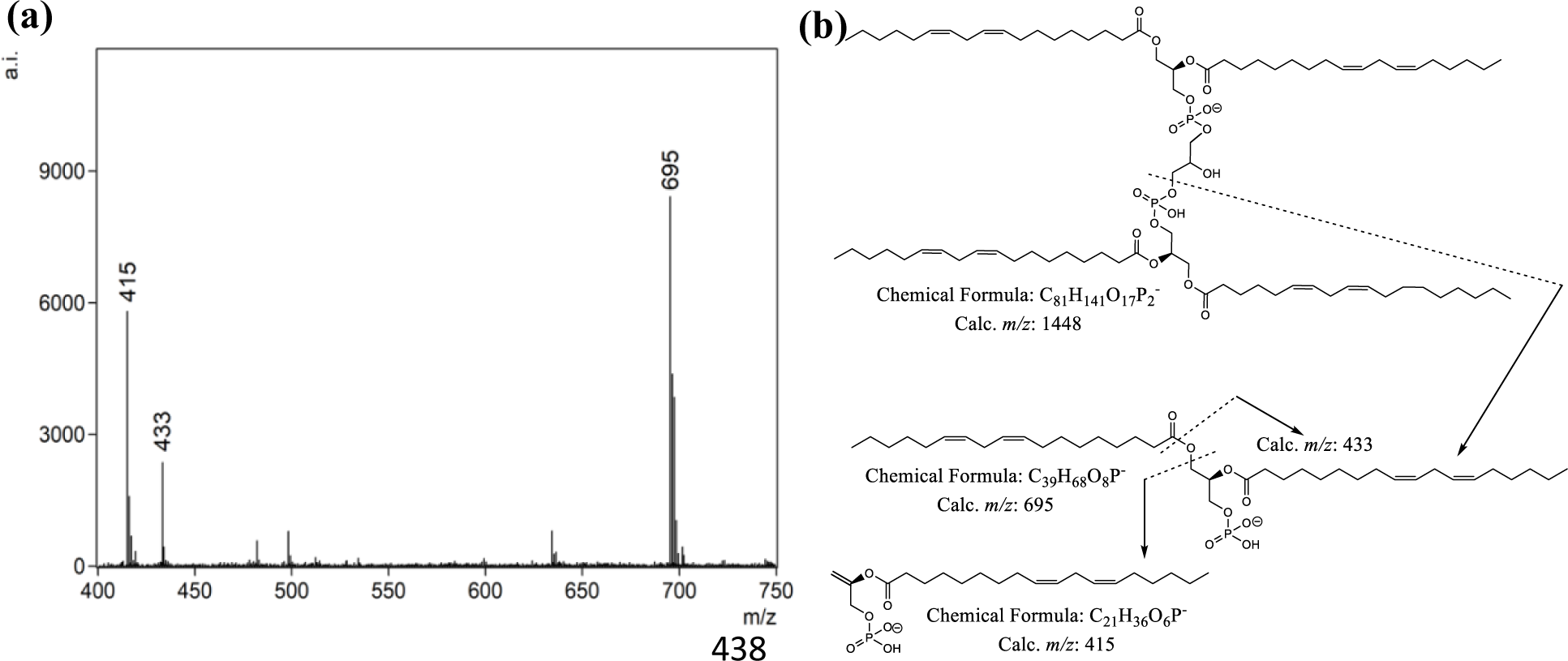
FLAT mass spectrum of 1448 interferences (a) A mass spectrum of *m/z* 1448 ion selected for tandem MS (b) Predicted structure of 1448 ion. All data from MALDI-timsTOF-MS.

**SI. Fig 2:**
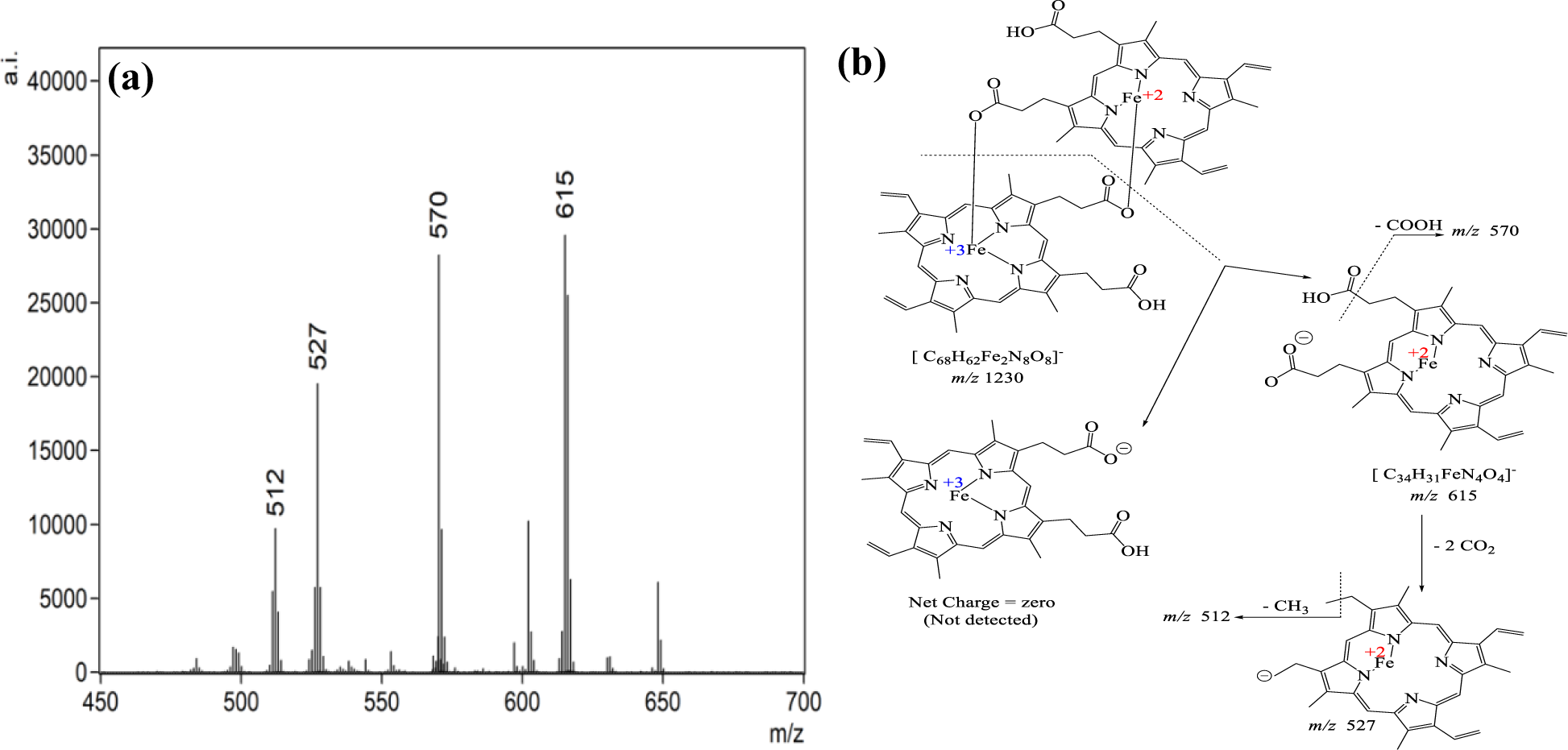
FLAT mass spectrum of heme with the FLAT analysis of urine samples with blood: (a) Tandem MS spectra of 1230 ion (b) Predicted structure of 1230 ion. All data from MALDI-timsTOF-MS.

